# The development of trait anxiety in nonhuman primates during the first year of life

**DOI:** 10.1101/2025.05.29.654506

**Authors:** Rachel Puralewski, Nakul Aggarwal, Jonathan Oler, Patrick Roseboom, Lauren Parkins, Marissa K Riedel, Carissa Boettcher, Ned Kalin

## Abstract

Anxious Temperament (AT) is a nonhuman primate phenotype that models childhood behavioral inhibition, a prominent risk factor for anxiety disorders and stress-related psychopathology. To understand its earliest antecedents, we characterized AT’s developmental trajectory over the first year of life. Infant monkeys (n=35, 24 female) were longitudinally phenotyped for AT by assessing threat-related increases in freezing behavior, reductions in coo-calling, and increases in circulating cortisol at 1.5, 6, 12, 24, and 52 weeks. AT levels increased linearly, reaching mature levels by 52 weeks. Individual differences in AT, and its components, were stable across development. These findings support further studies during early primate life aimed to uncover neurobiological factors mediating development of this phenotype, which in humans is linked to stress-related psychopathology.

## Introduction

Anxiety disorders (ADs) are common, typically emerge early in life, and are estimated to affect up to 20% of adolescents (Kessler et al., 2005; Rapee et al., 2023). Prior to onset and diagnosis of ADs, approximately 40-50% of children exhibit extreme behavioral inhibition (BI)(Chavira et al., 2002; Clauss & Blackford, 2012), which is a temperamental trait that is a significant risk factor for the later development of ADs. BI is characterized by marked shyness and extreme avoidance behaviors when children are confronted with potentially threatening situations, such as exposure to strangers or novelty (Kagan et al., 1987, 1989; Kagan & Snidman, 1999). To better understand how BI is associated with the development of pathological anxiety and ADs, it is important to characterize the earliest manifestations of BI and its maturational patterns during infancy and early childhood. This knowledge will lay the groundwork for efforts aimed at developing novel early-life interventions focused on increasing resilience in children at risk for developing ADs.

A nonhuman primate model using young rhesus monkeys, has been developed to better understand the neural mechanisms underlying extreme BI and the disposition to develop ADs (Kalin, 1993; Kalin & Shelton, 1989). Rhesus monkeys are ideally suited for translational research relevant to understanding developmental psychopathology, in particular ADs, as they share with humans similar threat-related emotional and behavioral responses (Ausderau et al., 2022), along with a prolonged period of parental rearing necessary for healthy psychosocial functioning and postnatal brain development (HARLOW & ZIMMERMANN, 1959; van der Horst et al., 2008). Importantly, rhesus monkeys and humans have similar cortical and subcortical brain structure and function as it relates to the expression of anxiety. In contrast to rodent species, rhesus monkeys have a well-developed prefrontal cortex, which plays an important role in integrating cognitive and emotional processes that are critical for the adaptive and maladaptive regulation of anxiety (Kenwood et al., 2022).

In children, BI is characterized by an individual’s tendency to inhibit ongoing behavior, remain still, and decrease speech when confronted with an unfamiliar person or novel situation (Kagan et al., 1987, 1988, 1989). The no-eye-contact condition (NEC) of the Human Intruder Paradigm (HIP) is used to assess this phenotype in young monkeys (Kalin & Shelton, 1989). The NEC entails separating a young monkey from its mother or conspecific, placing the subject in a test cage, and observing its behavioral responses to the presence of a human intruder standing 2.5 meters from the test cage while presenting their profile as an indirect threat to the monkey. It is critical that the intruder does not have eye contact with the monkey. The responses elicited by NEC are characterized by an overall inhibition of behavior. This involves decreases in locomotion and vocalizations, as well as freezing behavior, which is a complete cessation of movement and absence of vocalizations. Also, the NEC response is associated with an increase in HPA-axis activation as assessed by increases in plasma cortisol levels. Our work has expanded on the concept of BI by adding a physiological measure of stress reactivity, threat-induced plasma cortisol concentrations, to the phenotype, and we have termed this composite phenotype *anxious temperament* (AT) (Fox et al., 2008). Considerable work from our laboratory has demonstrated that individual differences in AT are stable over time, partially heritable, and associated with increased brain metabolism in a circuit that includes posterior orbitofrontal cortex, amygdala, bed nucleus of the stria terminalis, and the periaqueductal grey region of the brain stem(Fox et al., 2008, 2015; Kalin et al., 2005; Shackman et al., 2013).

Previous work with rhesus monkeys has focused on understanding AT and its neural correlates during the preadolescent period, which is a time of vulnerability in youth for the emergence of ADs. Gaining an understanding of the early developmental time course for the expression of AT is an important step towards investigating factors and mechanisms that lead to its extreme expression and risk to develop ADs. While some work has investigated the ontogeny of nonhuman primate threat responses, including components of AT (Kalin et al., 1991), this work used a cross-sectional design and examined human intruder responses at only a few timepoints during the first year of life. More specifically, this study (Kalin et al., 1991) found that around 2-4 months of age, infant rhesus monkeys begin to display adaptive threat-related responses (i.e., selectively expressing freezing behavior during NEC as compared to other types of threatening contexts that elicit active protective responses such as aggression (Kalin & Shelton, 1989)). This age roughly corresponds to 9-12 months of age in human children, which is the time at which stranger anxiety emerges (Brooker et al., 2013; Clauss & Blackford, 2012; Rheingold & Eckerman, 1973).

To build on these findings, we designed the current study using a within-subject longitudinal design with frequent AT phenotyping to advance the understanding of AT’s development during the first year of life. The goals of the study included tracing the longitudinal trajectories of the components of AT, characterizing individual differences in the development of AT, and assessing the stability of AT-related responses. Considerable work has identified the neural circuit underpinnings of AT in older, preadolescent, rhesus monkeys and in children with ADs and pathological anxiety (Birn et al., 2014a, 2014b; Fox et al., 2015, 2018; Shackman et al., 2013; Tromp et al., 2019). The findings from the current study will set the stage for understanding the development of these neural circuits as they relate to the earliest expressions of primate anxiety and the risk to develop ADs.

## Methods

### Study Design and Cohort Description

The study cohort consisted of 35 NHPs (*Macaca mulatta,* 24F, 11M). Animals were followed longitudinally across their first year of life, with each animal completing AT-related behavioral assessments at the same five ages between birth and one year (*see figure 1A*). This study also included longitudinal assessment of brain function and structure with multimodal brain imaging, with testing at similar timepoints as behavioral assessments discussed in this paper. As findings from neuroimaging data become available, they will be presented. Animals were housed with their birth mothers until 26 weeks, at which point they were weaned and pair-housed with a similarly aged peer from the same cohort. All monkeys were cared for in standard housing facilities at the Wisconsin National Primate Research Center in accordance with Institutional Animal Care and Use Committee protocols, including maintenance of 12-hour on/12-hour off light schedule, regular feeding, and age-appropriate enrichment. All behavioral data were collected during ‘lights-on’ period.

**Figure 1.**
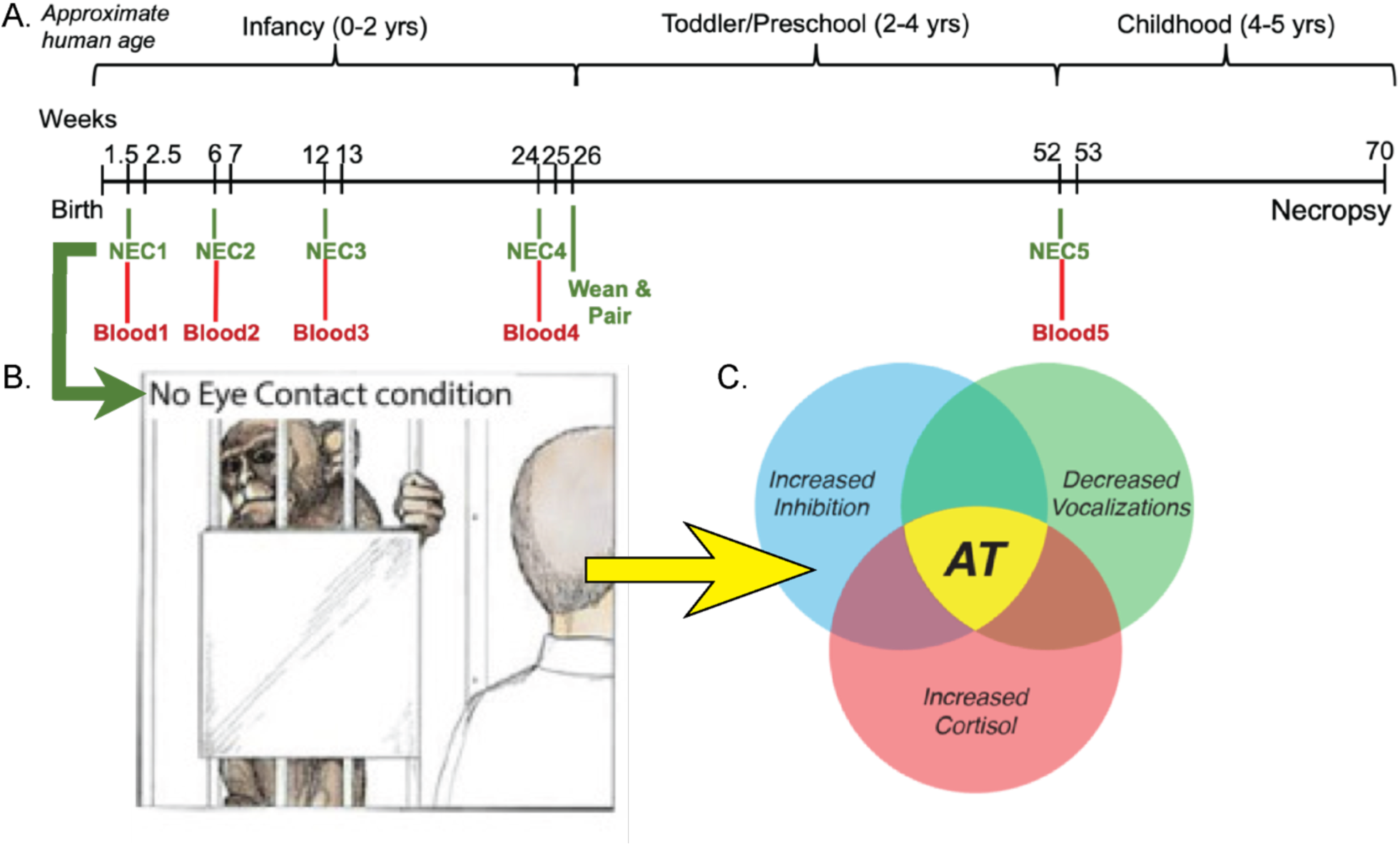
Overview of project timeline, behavioral paradigm, and Anxious Temperament, A) timeline of behavioral testing conducted on 35 NHP subjects (24 F, 11M). There were 5 timepoints of testing for each subject at 1.5, 6, 12, 24, and 52 weeks with 30 min NEC followed by collection of blood for cortisol assays, B) depiction of NEC paradigm, which typically elicits behavioral inhibition, recreated from original image depicted in “The Neurobiology of Fear (Kalin, 1993), C) AT is a composite score derived from behavioral expression during NEC (freezing duration and cooing frequency) and cortisol levels measured in blood collected following NEC, recreated from original image depicted in (Kalin, 2017).

Monkeys were each tested at five ages (∼1.5, 6, 12, 24, and 52 weeks) with 30 minutes of the NEC condition from the HIP (*see figure 1A*). At early ages, young study subjects were still clinging to, and thus housed with, their mothers on day of testing. At these early ages, mothers were first removed from the home cage and anesthetized with ketamine (15 mg/kg, IM). Then, infants were wrapped in a towel to keep warm and transported by hand to the testing room. Once older, pair-housed subjects were separated from their peer and moved to testing room via cage-transport. For NEC, monkeys were placed into a novel cage in an unfamiliar room. During the first 3 timepoints, a blanket-covered heating pad was placed on the cage bottom to ensure maintenance of infants’ body temperature during caregiver separation. A camera was positioned opposite the testing cage to capture behavior and vocalization sounds during the behavioral paradigm.

### No-Eye-Contact Exposure for evaluation of Anxious Temperament

Assessment of anxiety-related behavioral and neuroendocrine responses was performed through use of a modified version of the NEC. In this version, the monkey is placed into a novel testing environment (inner cage dimensions: 71 cm x 76 cm x 78 cm) and left alone momentarily to acclimate. Then, an unfamiliar intruder (a human whom the monkey has no contact with outside this context) enters the room. The same intruder was used for all 35 monkeys at all 5 ages. This intruder stood opposite the testing cage in view of the subject (approximately 2.5 meters from front of cage) with their profile facing the monkey for 30 minutes (Kalin et al., 2005; Kalin & Shelton, 1989) . Vocalizations and movement behaviors during NEC were scored using The Observer XT from Noldus (Zimmerman et al., 2009). Movements were scored as mutually exclusive duration behaviors; vocalizations were scored by vocalization type and frequency of occurrence. For this study, we specifically focused on duration of freezing, and frequency of coo vocalizations (coos) during the 30 min NEC. After NEC, monkeys were removed from testing cage, weighed, and anesthetized with ketamine (15 mg/kg, IM) under supervision of veterinary staff. During anesthesia, blood pressure, heart rate, and body temperature vitals were monitored. Once anesthetized, blood was drawn and spun down to separate out plasma. Plasma levels of the stress hormone cortisol (Cort) were assayed in duplicate using the commercially available ImmuChem coated tube radioimmunoassay (MP Biomedicals, Solon, OH). The intra-assay coefficient of variation (CV) was 4.9% and the inter-assay CV was 9.8%. The detection limit defined by the lowest standard was 1 µg/dL. Following sample collection, monkeys were monitored while anesthesia wore off and then were reunited with their mother or their cage-mate.

### Data Description and Statistical Analyses

Behavioral data was scored by four trained observers with high reliability between scorers (>90% agreement, kappa>0.8). Behavioral data was collected in bins of five minutes, and an average score per 5-minute bin was computed for each subject at each age. Vocalization frequency data was square-root transformed, and movement duration data was log_10_ transformed as is typical for these behaviors (Fox et al., 2008; Kalin et al., 2001; Kalin & Shelton, 1989). To account for diurnal cortisol rhythm, we computed residuals from models predicting cortisol blood levels from time of day (TOD) of sample collection. There was relatively little variability in TOD; all blood samples were collected between 8 AM-1 PM, with average collection time being 10:05 AM +/- 52 min (SD). Model residuals were used as cortisol values for all subsequent analyses.

All statistical analyses were conducted in R (version 4.2.3, https://www.R-project.org/). In formulating AT scores, transformed cortisol, freezing duration, and cooing frequency were first z-scored across all timepoints. As vocalization reduction is indicative of increased anxiety, cooing was combined with the other AT components as follows:

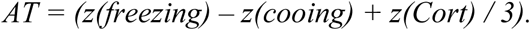

#### Stability Analyses

Stability of *between*-subjects’ individual differences for each measure across timepoints was examined using Pearson correlations, and p-values were determined with individual linear models predicting each timepoint from each other timepoint. We used the Bonferroni method to correct for multiple comparisons; with 10 between-timepoint pairwise comparisons, our adjusted threshold for significance is p<0.005. We conducted intraclass correlation (ICC) analyses to evaluate stability of measurements *within* subjects across the developmental timepoints. For our ICCs, we chose average two-way mixed effects models for consistency (ICC(3,k).

#### Determination of Developmental Trajectories

Individual developmental trajectories for variables of interest (behavioral, neuroendocrine, AT composite) were determined using within-subject linear mixed-effects models controlling for gestation length and sex, and including a random intercept to account for magnitude differences across subjects. Logarithmic growth patterns with increasing age were tested within the linear model framework by using log_10_ transformed age values. Linear, quadratic, & cubic shapes for the age-associated changes were also tested, as well as an *a priori* model testing a unique slope between each pair of timepoints. Growth trajectory model fits were assessed by comparing Bayesian Information Criteria (BIC) values, with the lowest (>2) BIC value indicating a model with the best fit. In cases where models had BIC values within 2 points, the simpler model was chosen. Models including random slopes, in addition to a random intercept, were also tested. We did not find that the inclusion of random slopes (for any variable of interest) improved the fit of the model (did not lower BIC values), nor was the random slope variable found to be significant within the best fit model. Therefore, we concluded that any differences between subjects for our variables of interest were likely associated with differences in magnitude and not differences in the rate of change. Multiple comparison correction was performed using the Bonferroni method. With 4 variables of interest, our adjusted p-value for developmental trajectories was p=0.0125. For all variables of interest, we performed pairwise t-test comparisons between all timepoints to evaluate if magnitude differences between timepoints were statistically significant. Using the Bonferroni method, our corrected p-value threshold of significance with 10 comparisons for each variable was p<0.005.

#### Statistical Comparisons to Mature Cohort

We used the Wilcoxon Rank Sum Test to investigate how distributions of AT scores from the current cohort compared with a larger cohort of 721 preadolescent naïve monkeys accumulated from prior studies in the lab that included 30 minutes of NEC, in addition to pairwise tests that compared how different timepoints from the current study compared to one another. Given that AT scores are computed relative to others in a population, AT was calculated within the data sample being used for the comparison. For details on analytical comparisons with the larger preadolescent monkey cohort, see supplement.

## Results

### Longitudinal trajectories of Anxious Temperament and its components across the first year of life

#### NEC-induced Freezing

Over the first year of life, infant monkeys demonstrated a 5-fold increase in freezing behavior from 1.5 weeks to 1 year of age (1.5 weeks average: 60 sec; 1 year average: 313 sec). A cubic equation was determined as the best fit for the within-subject age trajectory of freezing duration (*See Table 1 and* Figure 2A, p<2.2×10^-16^). Using within-subject linear modeling, we observed a decrease in freezing from 1.5 to 12 weeks, with the levels of freezing at 6 and 12 weeks significantly lower than freezing levels at 1.5 weeks (p<6.3×10^-4^ , p<5.3×10^-4^ respectively, see *Table 3*). Then, NEC-related freezing duration increased, with individuals having significantly greater freezing duration at 6 and 12 months compared to 12 weeks (p<2.98×10^-7^, p<2.1×10^-16^ respectively, see *Table 3*). Levels of freezing at 12 months were significantly greater than all previous ages (p<5.07×10^-11^).

**Table 1.**
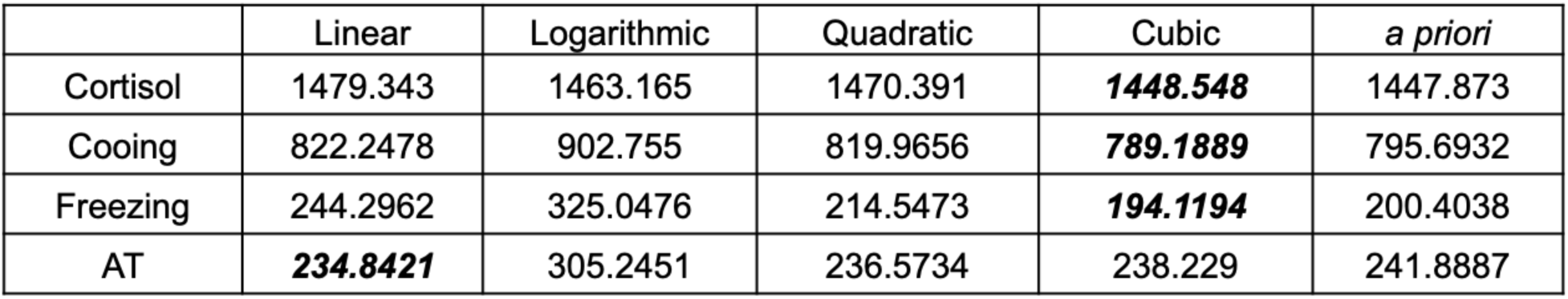
Bayesian Information Criteria (BIC) values for each shape tested for age-associated changes. The lowest BIC value (indicated in bold and italics) was used to select the best fit shape of age-related trajectory for AT and its components.

**Figure 2.**
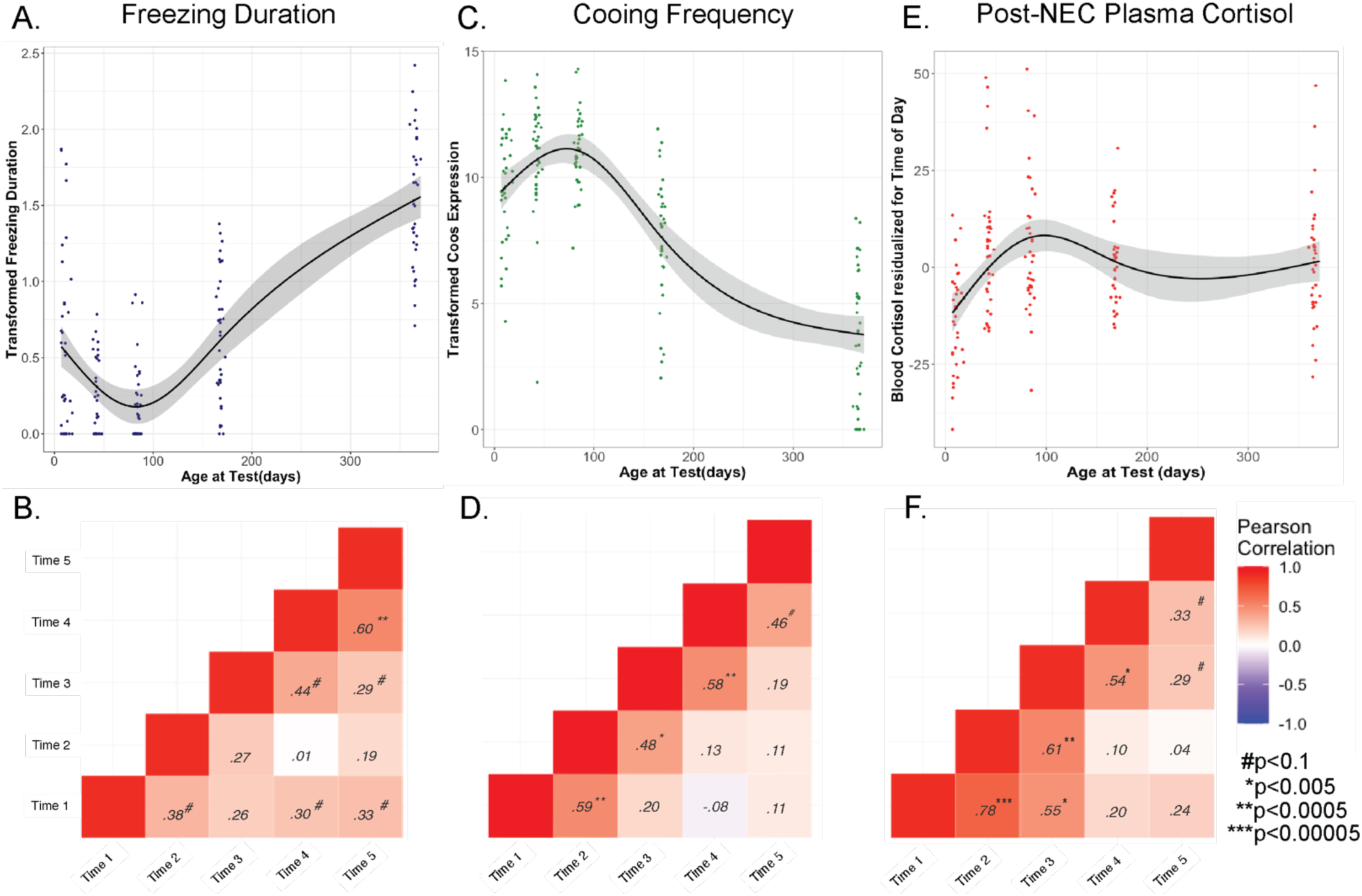
A) within-subject developmental trajectory of freezing (p<2×10^-16^), illustrating best fit of cubic age-related changes B) Pearson correlations examining freezing expression across timepoints, C) developmental trajectory of within-subject changes in cooing (p<2×10^-16^), depicting cubic age-related change; D) Pearson correlations examining cooing expression across timepoints, E) developmental trajectory of within-subject changes to cortisol (p<1.6×10^-9^), depicted with the best fit cubic age-related change F) Pearson correlations examining cortisol expression across timepoints.

Intraclass correlation (ICC) analyses across the 5 timepoints indicated “good” stability of freezing duration within individuals (*See Table 2*, ICC(3,k) = .67, p<3.2×10^-6^).

**Table 2.**
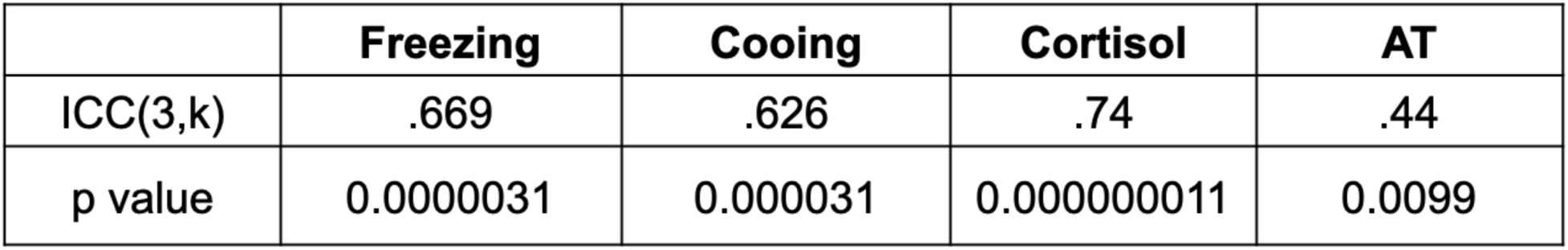
ICC(3,k) performance of measures across all timepoints, with corresponding p values.

Bivariate regression analyses across testing timepoints revealed moderate correlations (*r*’s ranging from .29-.60, p’s ranging from 0.09-1.3×10^-4^ *see* figure 2B). The correlation between freezing duration at ages 6 months and 1 year (r=.6, p<5.0×10^-4^) was significant, surviving multiple comparison correction. While not statistically significant after multiple comparison correction, it is noteworthy that freezing at 1.5 weeks predicted individual differences in freezing behavior at 1 year (*r*=.33, p=0.05).

#### NEC-induced Cooing Frequency

Across the first year of life, we observed a 4.75-fold decrease in cooing frequency (1.5 weeks average: 568.86 coos; 1 year average: 127.54 coos). A cubic fit was determined as the best equation for describing the age-related developmental trajectory of within-subject cooing frequency during NEC (*See table 1*, p<2×10^-16^, *and* figure 2C). From 1.5 to 12 weeks, monkeys showed an increase in NEC-related cooing (6 weeks comparison to 1.5 weeks: p<3.3×10^-4^, 12 weeks comparison to 1.5 weeks: p<3.8×10^-4^, see *Table 3*). From 12 weeks to 6 months, NEC-induced cooing decreased (p<0.002, see *Table 3*), which continued up to the 1-year assessment point. At 1 year of age, cooing frequency was significantly lower than all prior ages of testing (p’s<1.0×10^-10^, see *Table 3*). ICC(3,k) performance for cooing across all 5 ages was considered “good” (see *table 2*, .63, p<3.2×10^-5^), indicating stability of individual differences during this developmental period. Bivariate regression analyses between timepoints revealed moderate to strong correlations (*r’*s ranging from .46-.59, p’s ranging 0.0055 – 0.0002 *see* figure 2D). We found significant correlations passing multiple comparison correction between ages 1.5 & 6 weeks, 6 & 12 weeks, and 6 months & 1 year.

**Table 3.**
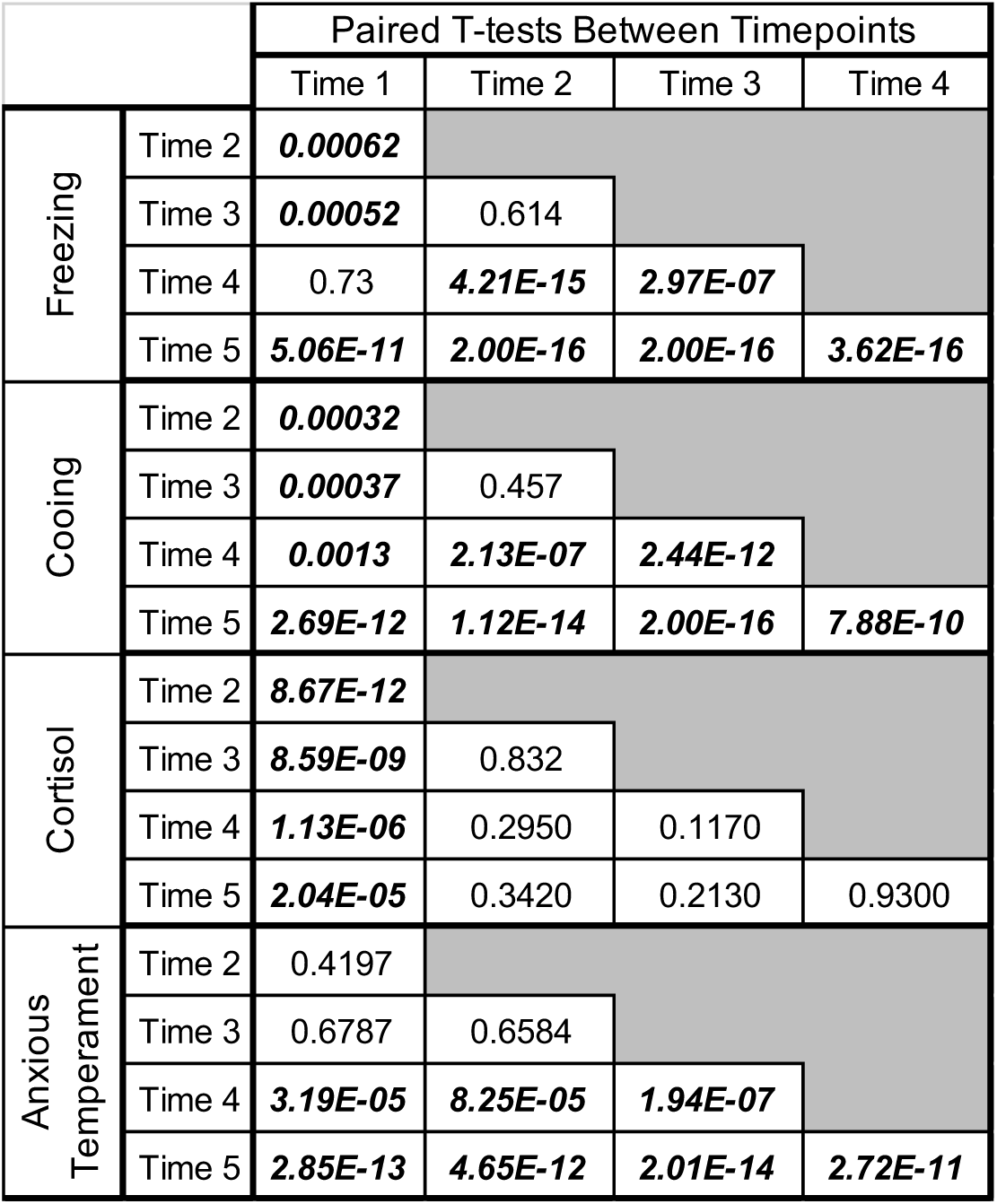
p-values for pairwise t-test comparisons across timepoints, comparisons with significantly different variable levels between timepoints indicated in bold italics.

#### NEC-related Plasma Cortisol

Threat-related plasma cortisol levels were 1.3-times greater at 1 year of age compared to 1.5 weeks (1.5 weeks average: 47.53 ng/mL; 1 year average: 62.35 ng/mL). We determined a cubic equation provided the best fit for the within-subject age-related developmental trajectory of NEC-related cortisol (See *Table 1*, *and* figure 2E, p<1.55×10^-9^). A significant within-subject increase in NEC-induced cortisol occurred from 1.5 to 12 weeks, with significantly greater cortisol levels at 6 and 12 weeks compared to 1.5 weeks (p’s<9×10^-9^). This relative increase was sustained through 6 months (6 month comparison to 1.5 weeks, p<1.14×10^-6,^), and 1 year (1 year comparison to 1.5 weeks, p<2.04×10^-5^). The ICC(3,k) for NEC-related plasma cortisol was considered “good” (.74, p<1.1×10^-8^, *see Table 2*). Bivariate regression analyses across adjacent timepoints revealed moderate to strong correlations (*r*’s: .33 - .78, p’s range: 0.054–4.8×10^-8^ *see* figure 2F), with the strongest associations being between the cortisol levels at 1.5 and 6 weeks. (*r* = .78, p<4.76×10^-8^). Significant correlations passing multiple comparison correction were found between ages 1.5 & 6 weeks, 1.5 & 12 weeks, 6 & 12 weeks, and 12 weeks & 6 months.

### Anxious Temperament across the first year of development

By combining the three above components, AT scores were computed. AT scores increased 2.5-fold between 1.5 weeks and 1 year of age. The best fit for within-subject age-related changes was linear (see *Table 1*) predicting a significant increase from birth to one year (p<2×10^-16^, *see* figure 3A). The monkeys’ AT levels at age 6 months and 1 year were significantly greater than all prior ages (p’s ranging 8.25×10^-5^ – 2.0×10^-14^). The levels of AT at 1 year were also significantly greater than the levels at 6 months (p<2.73×10^-11^). The AT levels at 1 year of age in this study represent mature AT levels based on comparison with data from a large sample (n=721) of rhesus monkeys phenotyped for AT, ranging from 9 months to 4.2 years of age (see supplement for details). ICC(3,k) analyses of AT across the first year of life indicated “moderate” stability (.44, p<0.0099, see *Table 2)* across all five ages. Bivariate regression analyses assessing adjacent ages demonstrated moderate to strong correlations (*r*’s ranging: .37-.62, p’s ranging 0.029-6.8×10^-6^ *see* figure 3B).

**Figure 3.**
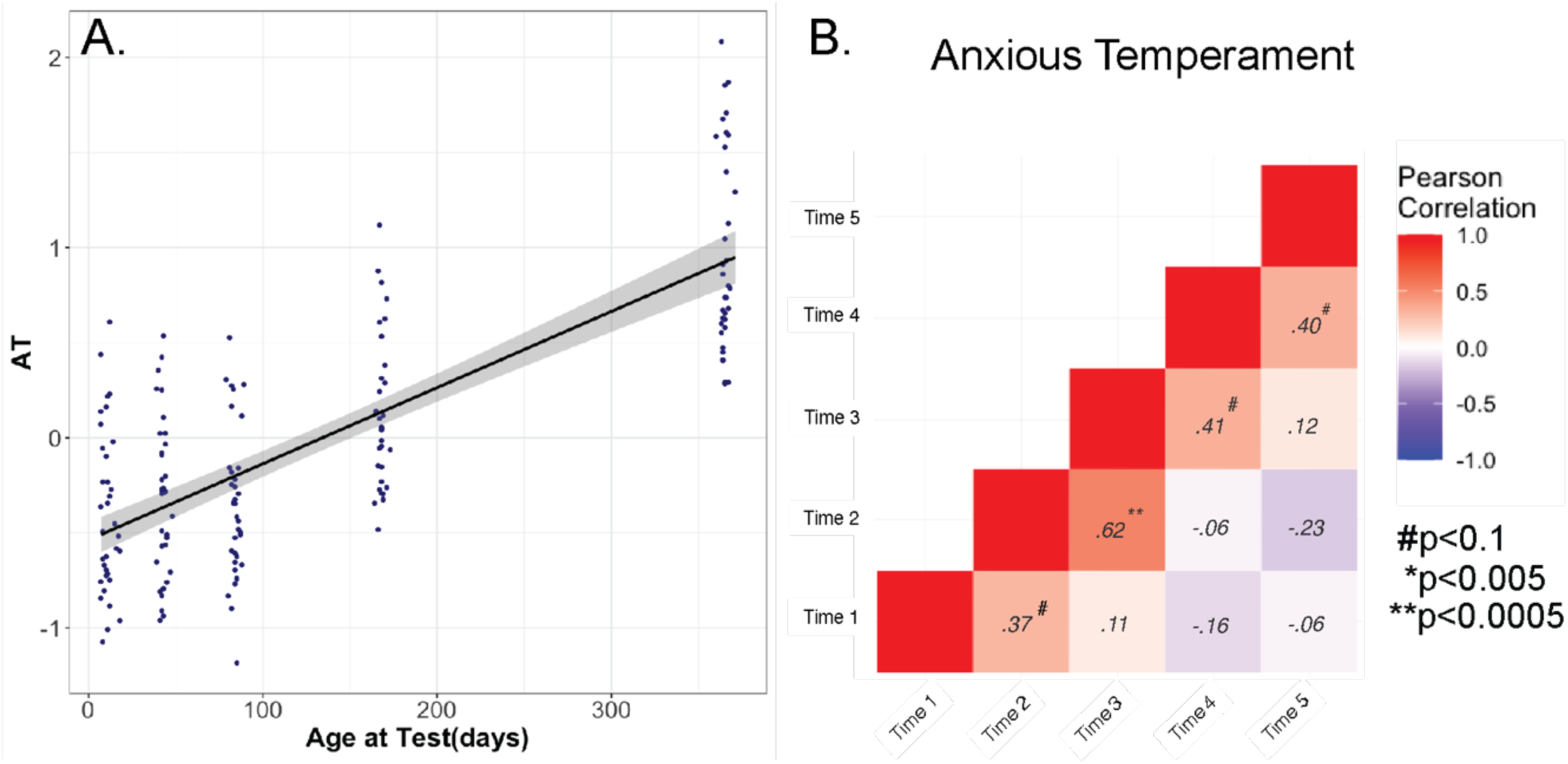
A) within-subject developmental trajectory of AT (p<2×10^-16^), with linear age-related change; B) pairwise correlations of AT across timepoints.

## Discussion

Behavioral inhibition (BI) is a childhood temperament known to confer significant risk for the development of anxiety disorders and stress-related psychopathology. The rhesus monkey AT phenotype (comprised of threat-related freezing duration, reductions in cooing frequency, and stress-related cortisol) is analogous to and extends the concept of human BI, and in this study, we demonstrate the early-life trajectories of AT and its components. As early as two weeks of age, infant rhesus monkeys display threat-related behavioral and hormonal responses that are the forerunners of mature AT. Over the course of the first 12 months of life, AT increases linearly and reaches mature levels by 12 months of age.

The adaptive response to NEC is to inhibit ongoing behaviors, including vocalizations, which serves to avoid detection by potential predators. Our data demonstrates that at the earliest ages after birth (1.5, 6, and 12 weeks of age), responses to NEC are characterized by relatively low levels of freezing accompanied by high levels of coo vocalizations. During infancy, high levels of coo vocalizations are indicative of separation distress, serving to enhance reunion with caregivers (Hinde & Spencer-Booth, 1967, 1971; Kalin & Shelton, 1989; Seay et al., 1962). When occurring at high levels during NEC early in life, it suggests that the drive for reunion with caregivers supersedes the self-preservational, inhibitory responses readily seen in more mature animals. At 12 weeks of age, the trajectories of NEC-induced freezing and cooing shift, indicative of a maturing, adaptive, inhibitory response to uncertain threat. Consistent with our finding that 12 weeks of age appears to be an inflection point in development, a previous cross-sectional study of rhesus monkeys (age 2-12 weeks) demonstrated that infant monkeys develop the capacity to adaptively regulate their behavioral responses to threat in the face of changing contexts at this age (Kalin et al., 1991). Others have found similar developmental trajectories for separation-induced vocalizations in early NHP life (Zhang et al., 2012). Taken together with our current data, it is likely that these developmental shifts in response to NEC reflect a transformation of the infant monkey’s ability to more independently cope with threat.

The developmental trajectory of threat-related cortisol demonstrates a relative stress hyporesponsive period when assessed at 1.5 weeks of age, which reaches mature levels by 6 weeks. This early HPA-axis hypo-responsiveness to threat is consistent with data from rodent and human studies (Gunnar et al., 2015; Gunnar & Donzella, 2002; Gunnar & Hostinar, 2015; Jansen et al., 2010; Levine, 2001; Rosenfeld et al., 1992). This phenomenon has been less well defined in rhesus macaques; a number of studies demonstrate HPA-axis activation in young rhesus monkeys (ages ranging from 2-9 months) when exposed to a stressor in the absence of their mother (Clarke, 1993; Koch et al., 2014; Levine et al., 1985; McCormack et al., 2009; Sánchez et al., 2005; SUAREZ-JIMENEZ et al., 2013; Thomas et al., 1995). We are aware of no studies examining HPA-axis reactivity in rhesus macaques during the first weeks of life, and in the presence of the indirect threat of an unfamiliar stranger. Our work provides information that fills this critical gap in the literature, supporting a period of HPA-axis hypo-responsiveness to indirect threats during the earliest phases of infant primate development. It has been speculated that the cortisol hyporesponsive period may function to protect the developing brain from high levels of glucocorticoids (Gunnar & Donzella, 2002; Gunnar & Hostinar, 2015; Herzberg & Gunnar, 2020).

Despite the magnitude of changes in AT and its components across development, the ICC analyses indicate stability of AT and its components within subjects, indicative of the trait-like nature of these measures that is evident early in life. Bivariate regression analyses assessing the relation between adjacent ages provides complementary evidence regarding the stability of these measures and their trait-like quality across this developmental period. Notably, freezing behaviors at the earliest timepoint (∼1 week) were moderately predictive of individual differences in freezing at 1 year of age. Taken together these data suggest that individual differences in traits that are components of AT can be reliably identified during the earliest weeks of a primate’s life. This is especially interesting in relation to children’s risk to develop ADs (Biederman et al., 1993; Kagan & Snidman, 1999), extending the potential period of identifying risk from childhood to early infancy.

In summary, we present the developmental time course for the emergence and maturation of AT and its threat-related behavioral and hormonal components in infant rhesus monkey. We observed robust increases in AT, associated with increases in threat-related freezing and decreases in threat-related cooing as monkeys developed during their first year. Consistent with an HPA-axis stress hypo-responsive period, we also observed reduced threat-related cortisol at 1.5 weeks, which increased at 6 weeks of age and remained at this higher level from 6 weeks to 1 year of age. Despite robust changes in the levels of AT and its components across this developmental period, we found individual differences in these parameters to be moderately stable across the first year of life. The stability and predictive natures of translational nonhuman primate threat-related responses almost immediately after birth resembles findings in human infancy (Brooker et al., 2013; Morales et al., 2022; Niermann et al., 2019; Planalp & Goldsmith, 2020; Tang et al., 2020), highlighting the potential utility of early-life phenotyping in relation to understanding the later development of trait-like individual differences associated with the risk to develop psychopathology. Future work aimed at characterizing the factors, during the first year of primate life, that contribute to the individual developmental patterns of AT will be important in understanding mechanisms that increase the early-life risk for individuals to develop ADs and other stress-related psychopathology.

## Supporting information

Supplemental Methods and Results

## Notes

### Competing Interest Statement

We report the following potential conflicts of interest. Ned Kalin receives research grants from the National Institutes of Mental Health; serves as a consultant to Pritzker Neuropsychiatric Disorders Research Consortium, Skyland Trail Advisory Board, the Early Adversity Research External Scientific Advisory Board at the University of Texas at Austin, Corcept Therapeutics Incorporated, Galen Mental Health Scientific Advisory Board, and EMA Wellness Scientific Advisory Board; and is the Editor in Chief for the American Journal of Psychiatry. No other authors have any potential conflicts of interest to report

### Funding

- National Institutes of Health: P51-RR000167
- National Institute of Mental Health: P50 MH100031
- Office of the Director: P51-OD011106

