## Supplemental Methods and Results for "The development of trait anxiety in nonhuman primates during the first year of life"

Supplement

### Supplemental Methods

*Cohort Description for older preadolescent data set*

Using a large cohort of 721 previously characterized preadolescent animals with ages from 9 months to 4.2 years (n=721, 386 M/ 335 F; 11 subjects < 1 yrs old, 443 subjects between 1-2 yrs, 191 subjects between 2-3 yrs, 72 subjects between 3-4 yrs, 4 subjects > 4 yrs), we sought to determine the age at which young rhesus monkeys express mature threat-related responses. Anxious Temperament (AT) was computed within this cohort by combining freezing duration, cooing frequency, and plasma cortisol levels as described in the methods section from the main article text. We investigated age-related effects within this cohort using linear mixed effects models predicting AT from age. We further assessed maturity within the sample, through use of the Wilcoxon Rank Sum test for comparison of the AT distributions of the youngest animals (n=113 subjects <1.25 yrs) to the eldest animals (n=271 animals >2 yrs) within this cohort. We used the Wilcoxon Rank Sum test to compare distributions of each age from the study sample (n=35) to the mature naïve sample of preadolescent monkeys (n=721). For these analyses, AT was calculated within each group for comparison. With 5 comparisons, we considered distributions to be significantly different from one another at a threshold of $p<0.01$.

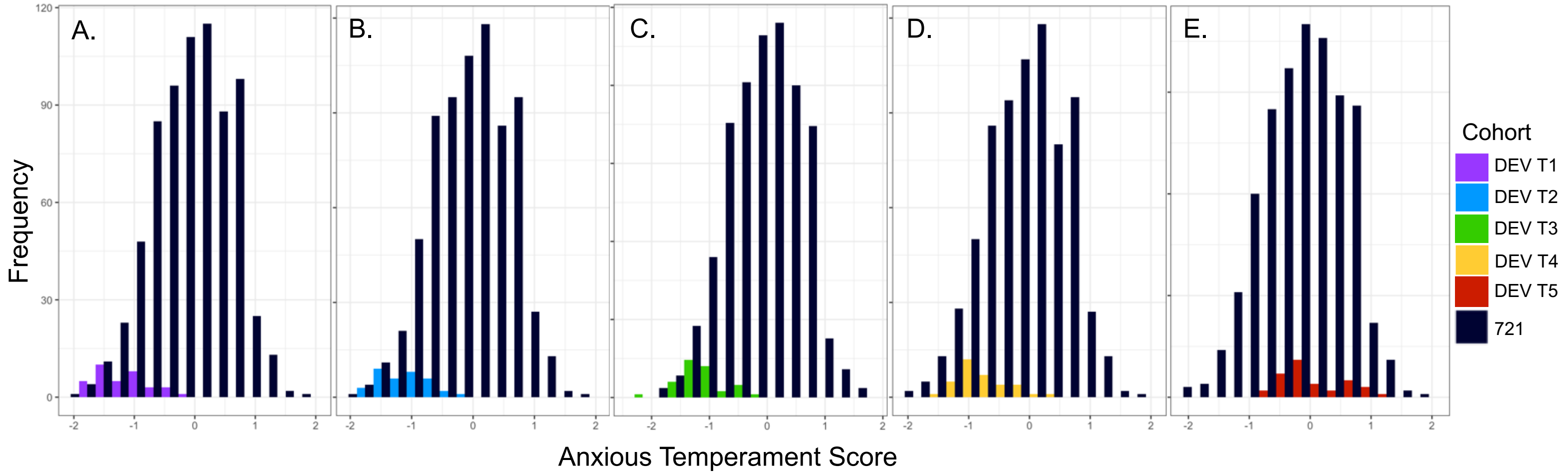


*Supplemental Figure S1 – A)AT scores at T1(age 1.5 weeks) have significantly different distribution compared to distribution of mature 721 NHP cohort AT scores, $p<2.0x10^{-16}$ ; B)AT scores at T2 (age 6 weeks) have significantly different distribution compared to distribution of mature 721 NHP cohort AT scores, $p<2.0x10^{-16}$ ;C)AT scores at T3 (age 12 weeks) have significantly different distribution compared to distribution of mature 721 NHP cohort AT scores, $p<2.0x10^{-16}$ ; D) AT scores at T4 (age 6 months) have significantly different distribution compared to the distribution of mature 721 NHP cohort AT scores, $p<4.6x10^{-14}$ ;E) distribution of AT scores at T5 (age 1 year) are not significantly different when compared to distribution of mature 721 NHP cohort AT scores, p = 0.3798.*

### Supplemental results

*Are AT levels at 1 year representative of mature AT levels ?*

Using a large cohort of 721 previously characterized preadolescent animals (ages 9 months to 4.2 years), we sought to determine the age at which young rhesus monkeys express mature threat-related responses. We first investigated any effects of age within this n=721 cohort, and did not find a significant age effect on AT ($p > 0.1$). To further evaluate maturity within this sample, we examined how the distribution of AT scores in the youngest animals (ages 9-15 months, n = 113) compared to the distribution of AT scores for the most matured animals (ages 2 – 4.2 years, n = 271). We found AT scores did not significantly differ in their distributions ($p = 0.212$) between these two age groups, demonstrating that AT levels around 1 year are comparable to those in older monkeys.

When examining how animals in our current study compare to the mature, preadolescent n=721 cohort, the Wilcoxon rank sum test demonstrated the distributions of AT scores at 1 year were not significantly different from the distribution of AT scores in the n=721 cohort (see *figure S1D*, $p=0.3798$), similar to what was found within the n=721 cohort. In contrast, the distribution of AT scores at 1.5 weeks, 6 weeks, 3 months & 6 months were significantly different from the distribution of AT scores measured in our n=721 population (see *Figure S1A, S1B, S1C, and S1D for T1, T2, T3, and T4* distribution comparisons, $p<1x10^{-13}$). Within our current developmental sample, we also tested whether the distribution of AT scores at 6 months were statistically different from those at 1 year, and found a significant difference ($p < 2.91x10^{-10}$). These results suggest that the AT phenotype reaches maturity sometime between 6 months to 1 year.
